# Towards Modular Engineering of Cell Signalling: Topographically-Textured Microparticles Induce Osteogenesis via Activation of Canonical Hedgehog Signalling

**DOI:** 10.1101/2023.07.19.549481

**Authors:** Fatmah I. Ghuloum, Lee A. Stevens, Colin A. Johnson, Natalia A. Riobo-Del Galdo, Mahetab H. Amer

## Abstract

Polymer microparticles possess great potential as functional building blocks for advanced bottom-up engineering of complex tissues. Tailoring the three-dimensional architectural features of culture substrates has been shown to induce osteogenesis in mesenchymal stem cells *in vitro*, but the molecular mechanisms underpinning this remain unclear. This study proposes a mechanism linking the activation of Hedgehog signalling to the osteoinductive effect of surface-engineered, topographically-textured polymeric microparticles. In this study, mesenchymal progenitor C3H10T1/2 cells were cultured on smooth and dimpled poly(D,L-lactide) microparticles to assess differences in viability, cellular morphology, proliferation, and expression of a range of Hedgehog signalling components and osteogenesis-related genes. Dimpled microparticles induced osteogenesis and activated the Hedgehog signalling pathway relative to smooth microparticles and 2D-cultured controls without the addition of exogenous biochemical factors. We observed upregulation of the osteogenesis markers *Runt-related transcription factor2* (*Runx2*) and *bone gamma-carboxyglutamate protein 2* (*Bglap2*), as well as the Hedgehog signalling components, *glioma associated oncogene homolog 1* (*Gli1*), *Patched1* (*Ptch1*), and *Smoothened* (*Smo*). Treatment with the Smo antagonist KAAD-cyclopamine confirmed the involvement of Smo in *Gli1* target gene activation, with a significant reduction in the expression of *Gli1*, *Runx2* and *Bglap2* (*p*≤0.001) following KAAD-cyclopamine treatment. Overall, our study demonstrates the role of the topographical microenvironment in the modulation of Hedgehog signalling, highlighting the potential for tailoring substrate topographical design to offer cell-instructive 3D microenvironments. Topographically-textured microparticles allow the modulation of Hedgehog signalling *in vitro* without adding exogenous biochemical agonists, thereby eliminating potential confounding artefacts in high-throughput drug screening applications.

**Graphical Abstract:** 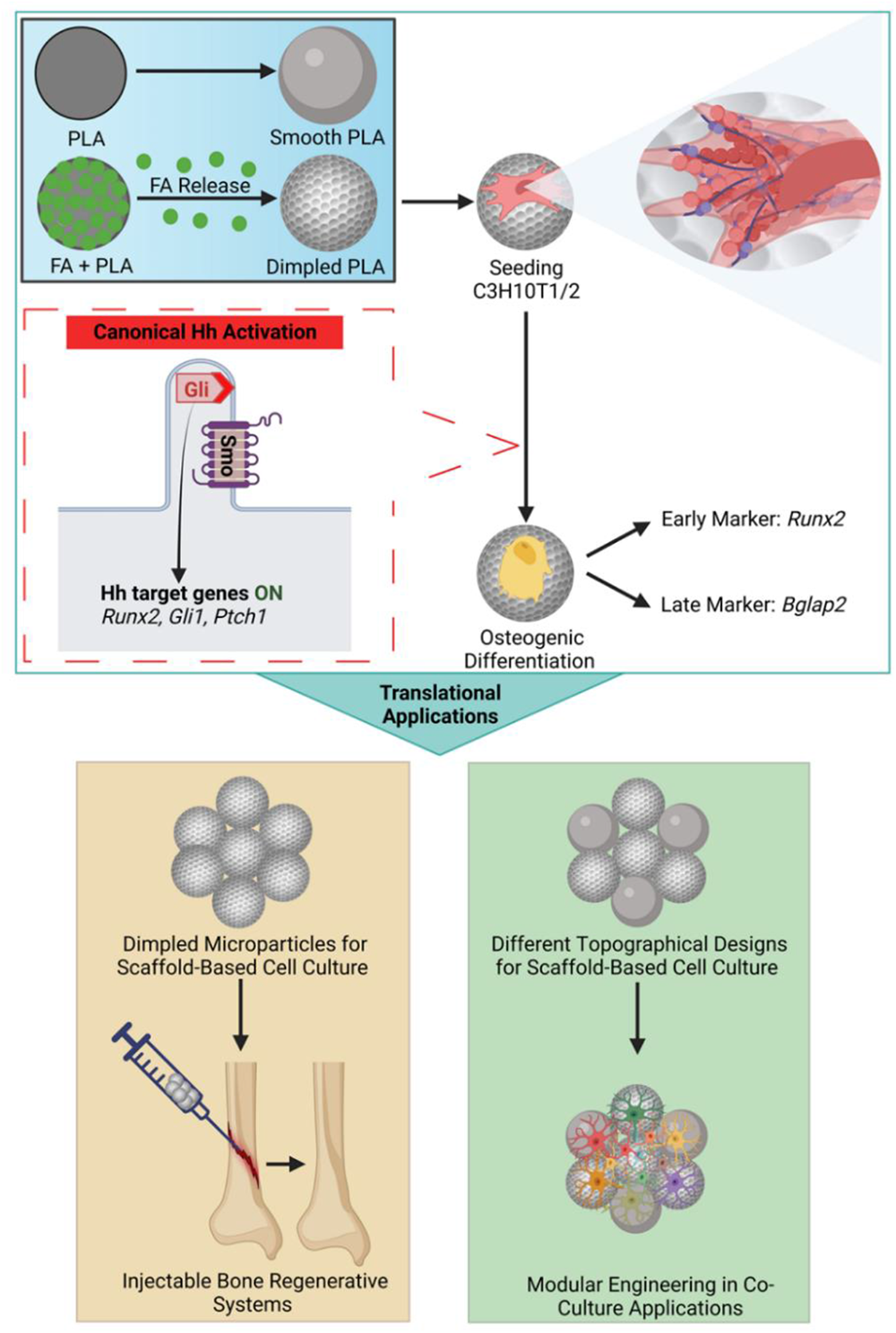

## INTRODUCTION

Significant advancements have occurred in the field of bioengineering in recent years, particularly with regards to cell culture strategies aimed at recreating the native cellular microenvironment and enhancing the relevance of *in vitro* experimental systems. This encompasses factors such as cell-extracellular matrix (ECM) interactions, biochemical signalling gradients, and mechano-transduction. A range of three-dimensional (3D) *in vitro* experimental systems have emerged in the last decade, ranging from organoids and hydrogel matrices, to engineering more complex 3D scaffold-based structures and microfluidics-based devices [1–3].

In the context of bone tissue engineering, tailoring these advanced cell culture experimental systems holds great promise. Native bone surfaces exhibit complex heterogeneous three-dimensional (3D) anisotropic topographies with irregular microstructures [4]. Mesenchymal stem cells (MSCs) can sense these topographical features and secrete distinct ECM components [5,6], which triggers mechanical responses at the cellular and sub-cellular levels in terms of cell morphology, migration, and tissue organisation [7]. Several studies have employed microscale topographical features to modulate MSCs differentiation in vitro, demonstrating that surface topographies can enhance the osteogenic differentiation of MSCs in the presence of classical osteogenic stimuli such as differentiation media or hydroxyapatite [8–10].

Microparticles, also known as microcarriers, provide customisable 3D scaffolds that offer high surface area-to-volume ratio for cell expansion [11]. Whilst microparticles were originally applied for large-scale cell expansion in bioreactors, their applications have since expanded to include tissue engineering and drug delivery applications. We have reported that tailoring micron-sized textured topographies on the surface of polymeric microparticles can effectively induce the osteogenic differentiation of human MSCs (hMSCs) in the absence of exogenous biochemical additives [2]. Nonetheless, the molecular mechanisms underlying this topography-induced osteoinduction is unclear. Understanding the correlation between microparticles-based architectural cues, and associated cellular responses on the molecular level is critical to achieve predictable outputs for drug discovery and regenerative medicine applications.

The Hedgehog (Hh) signalling pathway is a key mechanoresponsive signalling pathway that plays an essential role in bone regeneration by promoting osteoblast proliferation, differentiation and maturation [12,13]. Canonical Hh signal transduction in vertebrates involves Hh ligands binding to Patched (Ptch1), leading to Smoothened (Smo) activation and translocation to the primary cilium, where it prevents the processing and stimulates full activation of the Gli family transcription factors. This results in the transcription of Hh-target genes, including Gli1, Ptch1, and Runt-related transcription factor 2 (Runx2) [14]. Several studies have demonstrated that restoring the activity of the Hh signalling pathway in patients with bone fractures promotes bone regeneration [15–17]. In addition, dysregulation of *Gli1* expression, which normally promotes osteoblast differentiation and represses osteoblast maturation to maintain normal bone homeostasis, leads to a decrease in bone mass and a delay in fracture healing [18].

The mechanical features of culture substrates have been reported to play a key role in modulating cellular responses, including the expression of Hh-related genes [19–21]. Employing biphasic calcium phosphate scaffolds with rough micro- and macro-topographical features significantly increased the expression levels of Gli1 and osteoblast-related genes in human MG63 osteoblast-like cells compared to controls [22]. Lin *et al.* reported that topographical features of titanium surfaces can induce osteogenic differentiation of MG63 cells by upregulating the expressions levels of Hh-related genes, such as Gli1 and Smo, compared to smooth surfaces [23].

Surface engineering of microparticles to create tailored, cell-instructive substrates and direct downstream signalling effectors offers the opportunity to transform cell culture substrates and cell delivery systems from passive components to tailored, functional ones. This offers a powerful tool for developing advanced experimental systems to generate novel biological insights, and a potential alternative to conventional biochemical supplements that can have potential pleiotropic effects on signalling cascades and trigger off-target cellular responses in tissue engineering applications. This study focused on uncovering the role of the Hh–Gli signalling activation in the topographically-induced osteogenesis of mesenchymal progenitors observed on topographically textured microparticles [2].

## METHODS

### Fabrication of smooth and dimpled microparticles

Poly(D,L-lactic acid) (PLA) microparticles (Ashland Viatel DL 09 E; Mn 56.5 kDa, Mw 111 kDa, 1, IV 0.8-1.0 dl/g, Ashland Specialties UK) were prepared by a solvent evaporation oil-in-water emulsion technique as described previously [2]. The organic phase, containing 1g of PLA in dichloromethane (20% (*w/v)* in DCM; Thermo Fisher Scientific) was homogenised (Silverson Machines Ltd, UK) at 3800 rpm for 5min in an aqueous continuous phase, containing 1% (*w/v)* of poly(vinyl acetate-co-alcohol) (PVA; MW 13–23 kDa, 98% hydrolysed; Sigma-Aldrich) as stabiliser. The resulting emulsion was stirred continuously at 480 rpm for a minimum of 4 h. Microparticles were then centrifuged at 3500*xg* for 5 min and subsequently washed with deionised (DI) water. Microparticles of size range 40-70 µm were collected and subsequently freeze-dried.

To produce dimpled microparticles, 23% (*w/w*) fusidic acid (FA; 98%, 5552333, Thermo Scientific Acros) in PLA was incorporated (FA/PLA total content of 10% *w/v*) at 1200 rpm into PVA solution as previously described [2]. Dimpled microparticles were obtained after FA release in phosphate buffered saline (PBS; 0.1 M, pH 7.4, 0.15 M NaCl, Gibco) at 37°C for 7 days.

### Particle size analysis

Samples were dispersed in deionised water, and laser diffraction was used to measure microparticle sizes using a Camsizer XT (Retsch Technology, Germany) for 5min. For each batch of microparticles, polydispersity index (PDI) was used as a measure of broadness of size distribution. PDI of the fabricated microparticles was defined as the square of the standard deviation divided by their mean diameter. Dimples generated were characterised using ImageJ software by measuring the diameter of a minimum of 250 dimpled microparticles over five independent SEM images for two fabricated batches.

### Brunauer-Emmett-Teller (BET) surface area measurements

Surface area of the fabricated microparticles were calculated as detailed previously [2]. Krypton (Kr) sorption isotherms were carried out using a Micromeritics ASAP 2420 (Micromeritics, Norcross, GA, USA) at -196°C. Approximately 500 mg samples were degassed under high vacuum (<10 mtorr) at 37°C for 48 h to remove moisture and other adsorbed gases. As Kr solidifies at higher pressures at -196°C (>1.77 torr) [24], sorption isotherms were carried out from 0.10 to 0.65 relative pressure. The specific surface area was acquired from 0.05-0.30 relative pressure using the BET model. The pore volume/size distribution was extrapolated from the entire adsorption isotherm (1.75-20 nm) using the Derjaguin–Broekhoff–de Boer model [25].

### Cell culture

C3H10T1/2 cells (ATCC-CCL-226) were kindly gifted by Dr. Hazel Fermor. C3H10T1/2 cells were maintained in Dulbecco’s modified Eagle’s culture medium (DMEM; 21969-035, Gibco), which was supplemented with 1% (*w/v*) L-Glutamine (Gibco), 1% (*w/v*) Penicillin-Streptomycin (Gibco) and either with 10% (*v/v*) fetal bovine serum (FBS) (Gibco) “regular media” or 2% FBS “serum-reduced media”, at 37°C with 5% carbon dioxide (CO2) atmosphere in a humidified incubator. StemPro™ Differentiation Kits (Gibco, UK) were used to confirm tri-lineage differentiation potential of the cells.

### Preparation of microparticles for cell culture

Microparticles were placed in cell-repellent CELLSTAR® 96 well plates (Greiner bio-one) and sterilised with UV light at 254 nm for 30 min at 4x10^4^ mJ. The quantities of smooth and dimpled microparticles were calculated to obtain a surface area for cell attachment of 0.32 cm^2^ per well. Following this, microparticles were conditioned for 1h. Cells were seeded at 3x10^4^ cells/cm^2^ and placed on a plate shaker for 15 min for all conducted experiments. For 2D-cultured controls, cells were seeded at the same seeding density in tissue culture-treated 96 well plates (CytoOne, Thermo Fisher Scientific).

### Assessing cell viability and proliferation

To assess cell viability three days post-seeding, the Viability/Cytotoxicity Assay Kit for Animal Live & Dead Cells (Biotium, UK) was used according to the manufacturer’s protocol. Briefly, 1 μM calcein-acetoxymethyl (calcein-AM) and 4μM ethidium homodimer III (EthD) were added per well. Cells were then imaged with an EVOS FL Auto 2 imaging system (Thermo Fisher Scientific) using FITC and TexasRed^®^ filters, respectively.

Proliferation was assessed using PrestoBlue™ Cell Viability Reagent (Invitrogen, UK) at 1-, 3-and 10 days after seeding according to the manufacturer’s protocol. Briefly, culture medium was replaced by PrestoBlue™: culture medium (1:9) and incubated for 1h at 37 °C. After incubation, 100 µL of Prestoblue solution from each well was measured at λ_exc_/λ_em_ 560/590nmusing a microplate reader (Polarstar Optima, BMG Labtech).

### Scanning electron microscopy (SEM)

Samples were mounted onto double-sided copper tape (Agar Scientific) and placed on an aluminium pin stub (AGG301, Agar Scientific). Samples were sputter-coated with gold (Cressington sputter coater 208HR) for 4min prior to imaging, SEM images were taken using a Carl Zeiss EVO MA15 scanning electron microscope (Oxford Instruments) at 10-20 kV.

For sample preparation of cell-containing samples for SEM, cells were fixed three days post-seeding using 2.5% (*v/v*) glutaraldehyde (STBJ9947, Sigma-Aldrich). This was followed by sequential dehydration using ethanol (Fisher Chemical) at increasing concentrations (10, 25, 50, 80 and 100%). The fixed aggregates were transferred on aluminium pin stubs (AGG301, Agar Scientific) for SEM imaging. For measuring the average size of cellular protrusions observed on microparticles, three independent images were processed using ImageJ software.

### Immunocytochemistry

Cells were fixed using 3.7% (*v/v*) formaldehyde then permeabilised using 0.1% Triton X-100 (Alfa Aesar) for 30min. To block non-specific binding, incubation in 1% (*w/v*) bovine serum albumin (BSA, Sigma Aldrich) in PBS was used with either 10% (*v/v*) normal goat serum (G9023, Sigma-Aldrich) or normal donkey serum (D9663, Sigma-Aldrich), according to the species of the secondary antibody, for 1 h. Cells were then incubated with the primary antibody overnight at 4°C. The secondary antibody was added for 2 h the following day.

For F-actin staining, cells were stained with ActinGreen™ 488 ReadyProbes™ Reagent (AlexaFluor™ 488 phalloidin) (R37110, Invitrogen) and nuclei were counter-stained with NucBlue™ Fixed Cell ReadyProbes™ Reagent (R37606, Invitrogen). Cells were observed using a Zeiss LSM880 confocal laser scanning microscope (Carl Zeiss, Germany).

The following primary antibodies were used: anti-ADP-ribosylation factor-like protein (ARL13B) mouse monoclonal antibody (1:2000; clone N295B/66, NeuroMab, RRID: AB_2877361), human/mouse GLI-1 affinity purified polyclonal (1:100; AF3455, R&D systems, RRID: AB_2247710), RUNX2 (27-K) antibody (1:500; sc-101145, Santa Cruz Biotechnology, RRID: AB_1128251), and monoclonal anti-Vinculin (1:70; V4505, Sigma-Aldrich, RRID: AB_477617). Secondary antibodies used in this study were donkey anti-goat IgG (H+L) cross-adsorbed secondary antibody, conjugated to Alexa Fluor 594 (1:500; A-11058, Thermo Fisher Scientific, RRID: AB_2534105) and goat anti-mouse IgG (H+L) cross-adsorbed secondary antibody, conjugated to Alexa Fluor 647 (1:500; A-21235, Thermo Fisher Scientific, RRID: AB_2535804).

### Gene expression analysis using real-time polymerase chain reaction (RT-qPCR)

Total RNA was isolated using the RNAqueous™-Micro Kit (Thermo Fisher Scientific) following the manufacturer’s protocol. The concentration of RNA in each sample was measured using a NanoDrop™ One Microvolume UV-Vis Spectrophotometer (ThermoFisher Scientific). Reverse transcription of RNA was performed using iScript™ Select cDNA Synthesis Kit (Bio-Rad, USA) following the manufacturer’s protocol. T100 Thermal Cycler (Bio-Rad) was used following the reaction conditions listed in Table S1. Primer sequences are listed in Table S2. The transcript levels of *Gli1*, *Smo*, *Ptch1*, *Runx2*, osteocalcin (encoded by *Bglap2*), and Glyceraldehyde-3-Phosphate Dehydrogenase (*Gapdh)* were determined using SsoFast™ EvaGreen® Supermix (Bio-Rad), quantified using CFX96 C1000 thermocycler (Bio-Rad) and following the reaction conditions listed in Table S3. No template controls (NTC) for each primer set and no reverse-transcriptase controls (NRT) for each sample were run. Purmorphamine (Miltenyi Biotec) was used as a positive control for *Gli1, Smo,* and *Runx2* expression, dissolved in dimethyl sulfoxide (DMSO; 2µM) in serum-reduced media. Corresponding vehicle controls were prepared as 0.06% DMSO. Relative expression levels for each gene of interest compared to the housekeeping gene *Gapdh* were determined by using the 2^−ΔΔCt^ method [26] and normalised to the corresponding negative controls along with the standard deviation.

For measuring the expression of *Bglap2*, 2D-cultured C3H10T1/2 cells were treated with osteoinductive media containing dexamethasone (StemPro, Gibco, UK) as positive controls, or untreated as negative controls.

### Western Blotting

After 3 days in culture, cells were lysed using CelLytic™ M lysis buffer (C2978, Sigma-Aldrich) containing 1 mM phenylmethyl sulfonyl fluoride (PMSF) (36978, Thermo Fisher Scientific), 1 mM ethylenediaminetetraacetic acid (EDTA; Corning) at pH 8.0, and 1 µL protease inhibitor cocktail (P8340, Sigma-Aldrich). After incubation on ice for 20 min, the cell lysates were prepared by spinning down at 4°C, 13,000 rpm for 15 min. Total protein concentrations were determined using the Pierce™ Bicinchoninic acid (BCA) Protein Assay Kit (ThermoFisher Scientific, UK). Cell lysates were analysed by western blot with anti-Gli1 (L42B10) mouse monoclonal antibody (1:800, 2643, Cell Signaling Technology, RRID: AB_2294746) followed by goat anti-mouse IgG (H+L)-horseradish peroxidase (HRP)-linked secondary antibody (1:1000, 1721011, Bio-Rad, RRID: AB_11125936). Detection of the horseradish peroxidase signal was carried out using Clarity Western enhanced chemiluminescence (ECL) Substrate (Bio-Rad), and immunoreactive bands were visualised in a ChemiDocTM MP Imaging System using Image Lab software (Bio-Rad). Quantification of proteins was carried out by densitometry analysis using ImageJ software (v1.54b) and normalised to loading control HRP-Conjugated GAPDH (1:5000; HRP-60004, Proteintech, RRID: AB_2737588).

### Treatment with KAAD-cyclopamine

Prior to experiments, C3H10T1/2 cells were cultured in serum-reduced media for 24 h, then treated with 300 nM KAAD-cyclopamine (Abcam, UK) or 0.06% DMSO as vehicle control for 3 days before analysis.

### Statistical analysis

Statistical analysis was performed using GraphPad Prism version 9.3 (GraphPad Software Inc., San Diego, USA). Normality tests were performed, and data was analysed using unpaired Student’s *t*-test or parametric one-way/two-way Analysis of Variance (ANOVA) with Tukey or Dunnett’s *post-hoc* tests. All data is shown as the mean ± standard deviation (SD), with *p* ≤0.05 considered the threshold for statistical significance.

## RESULTS

### Fabrication and characterisation of topographically-textured microparticles

In this study, smooth and topographically-textured microparticles with specific surface micro-scale features were produced by exploiting the phase-separation of a sacrificial component using the solvent evaporation oil-in-water emulsion technique, as described previously [2] (Figure 1A-C).

**Figure 1:**
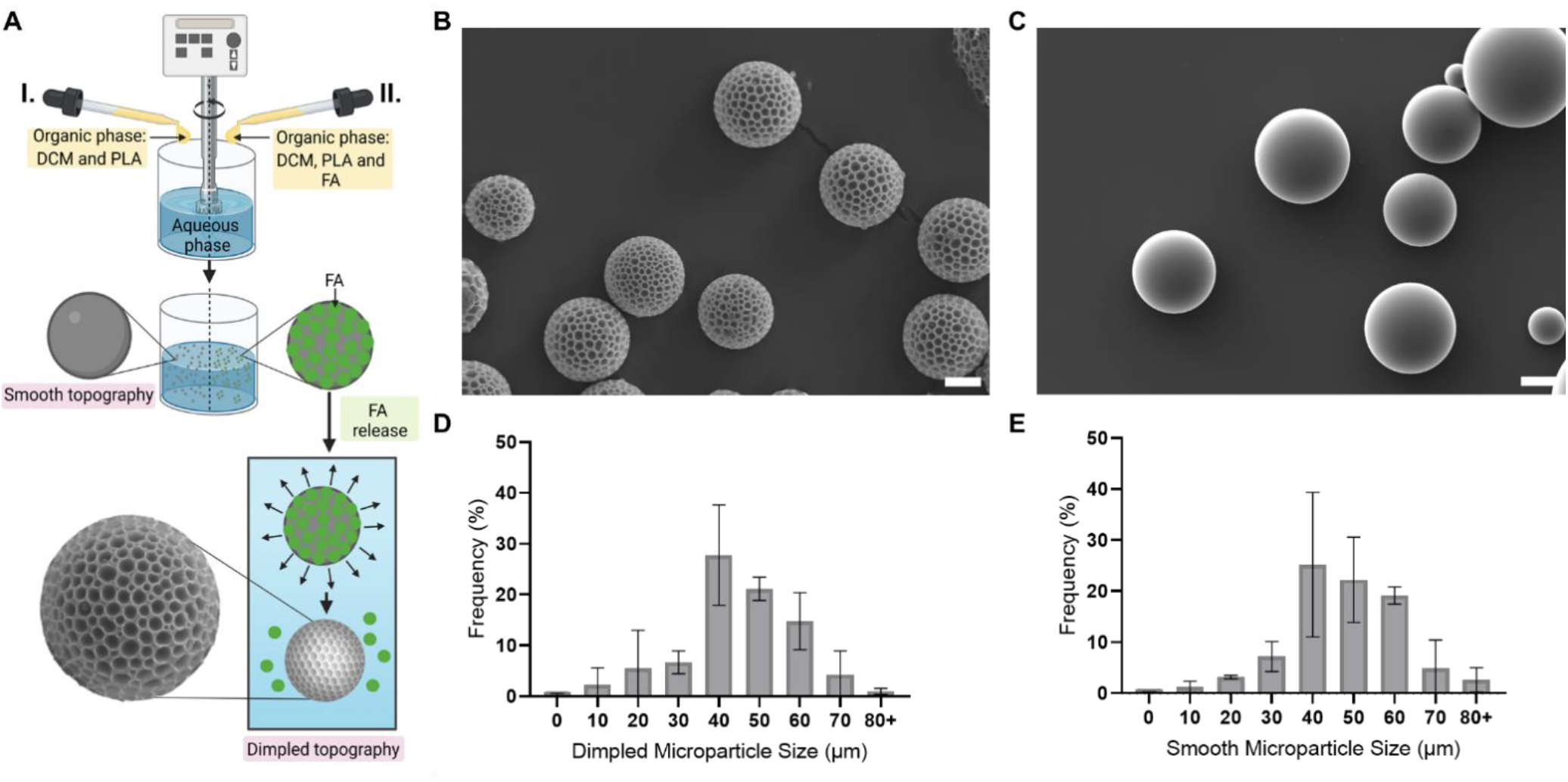
Fabrication of smooth and topographically-textured (dimpled) PLA microparticles. (A) Schematic representation showing the fabrication of smooth (I) and topographically-textured dimpled microparticles (II) by a modified oil-in-water solvent evaporation emulsion method. The topographical features are engineered using fusidic acid as a phase-separating sacrificial component. (B, C) Representative scanning electron microscopy images of dimpled (B) and smooth (C) microparticles, respectively, fabricated via a solvent evaporation oil-in-water emulsion technique (Scale bar= 20μm). (D, E) Graphs displaying the microparticle size frequency distribution for the batches used in this study. Abbreviations: PLA, Poly(D,L-lactic acid); DCM, Dichloromethane; FA, Fusidic acid.

Emulsion settings, such as homogenisation speed and polymer concentration, were optimised (Table 1) so that the fabricated smooth and dimpled microparticles were of a comparable mean diameter, in line with the reported sizes of osteo-inductive microparticles we previously demonstrated to induce osteogenesis in MSCs [2]. The mean diameter of smooth and dimpled microparticles was 53.21±14.18 μm and 50.15±14.16 µm, respectively (Figure 1D and 1E) with a mean dimple size of 4.6±0.27 μm on the surfaces of the dimpled microparticles (Table 1).

**Table 1:** Fabrication parameters and properties of microparticles used in this study.

|  |  | **Smooth** | **Dimpled** |
| --- | --- | --- | --- |
| **Emulsion settings** | FA/PLA ratio | 0/100 | 30/70 |
|  | Homogenization RPM | 3800 | 1500 |
|  | Polymer concentration (%) | 20 | 10 |
| **Particle properties** | Particle size ($\mu\text{m}$) | 53.21 ±14.18 | 50.15 ±14.16 |
| | Polydispersity index (PDI) | 0.08 ±0.04 | 0.07 ±0.02 |
| | Dimple size ($\mu\text{m}$) | - | 4.60 ±0.27 |
| | BET surface area ($\text{m}^2/\text{g}$) | 0.27 | 0.43 |
| | Vmicro ($\text{cm}^3/\text{g}$) | 0.0000096 | 0.0000175 |
| | Vmeso ($\text{cm}^3/\text{g}$) | 0.0001396 | 0.0002283 |

Abbreviations: FA, Fusidic acid; PLA, Poly(D,L-lactic acid); BET, Brunauer–Emmet–Teller; RPM, Rotation per minute; PDI, Polydispersity index; Vmicro, micropore volume; Vmeso, mesopore volume (we define mesopore volume as the pore volume originating from the pores smaller than 20 nm).

Surface areas (m^2^/g) of microparticles (Table 1) were calculated using the Brunauer-Emmett-Teller (BET) model. To ensure consistency of surface area available for cell attachment across samples, the mass of smooth and dimpled microparticles used was calculated, as previously described [2], to obtain a surface area of 0.32cm^2^ for consistent cell seeding across all 2D and 3D cultures. Both microparticle designs displayed negligible pore sizes, as reflected by their low volumes of micro-pores (Vmicro) and meso-pores (Vmeso) (Table 1).

### Effect of the Topographically-Textured Microparticles on Viability, Proliferation, and Morphology of C3H10T1/2 cells

C3H10T1/2 mouse-derived embryonic fibroblast cells have been widely utilised as a model of multi-potent mesenchymal cells [27,28]. The use of this standard cell line addresses the concern that selective cell attachment from heterogeneous populations of mesenchymal stromal cells may be responsible for the observed osteoinductive response when cultured on the different microparticle designs, rather than being attributable to active cellular differentiation. The ability of these cells to differentiate into osteogenic, adipogenic and chondrogenic lineages was confirmed (Figure S1).

To elucidate potential signalling mechanisms underpinning the osteo-inductive influence of topographically-textured microparticles [2], serum starvation was employed. Serum starvation is frequently used to synchronise all cells to the same cell cycle phase [29]. Furthermore, serum-starved cells terminate all serum-dependent signalling pathways, which eliminates background when studying the effect of external stimulation on cell behaviour [30,31]. Cells were cultured in low-serum conditions (2% *v/v* FBS) to allow for cell attachment and avoid stress that may trigger the activation of certain signalling pathways to promote cell survival [32]. To assess cell attachment and viability in the presence of these low serum conditions, cells were starved overnight before seeding on microparticles. The viability of mesenchymal progenitor C3H10T1/2 cells cultured on microparticles in serum-reduced media was compared to cells cultured in regular DMEM culture media supplemented with 10% FBS using the Viability/Cytotoxicity Assay after 3 days in culture. C3H10T1/2 cells successfully attached to the microparticles and showed high viability under both conditions (Figure 2A and 2B). Additionally, visibly smaller cell-microparticle aggregates were observed on dimpled microparticles compared to smooth microparticles in both media conditions.

**Figure 2:**
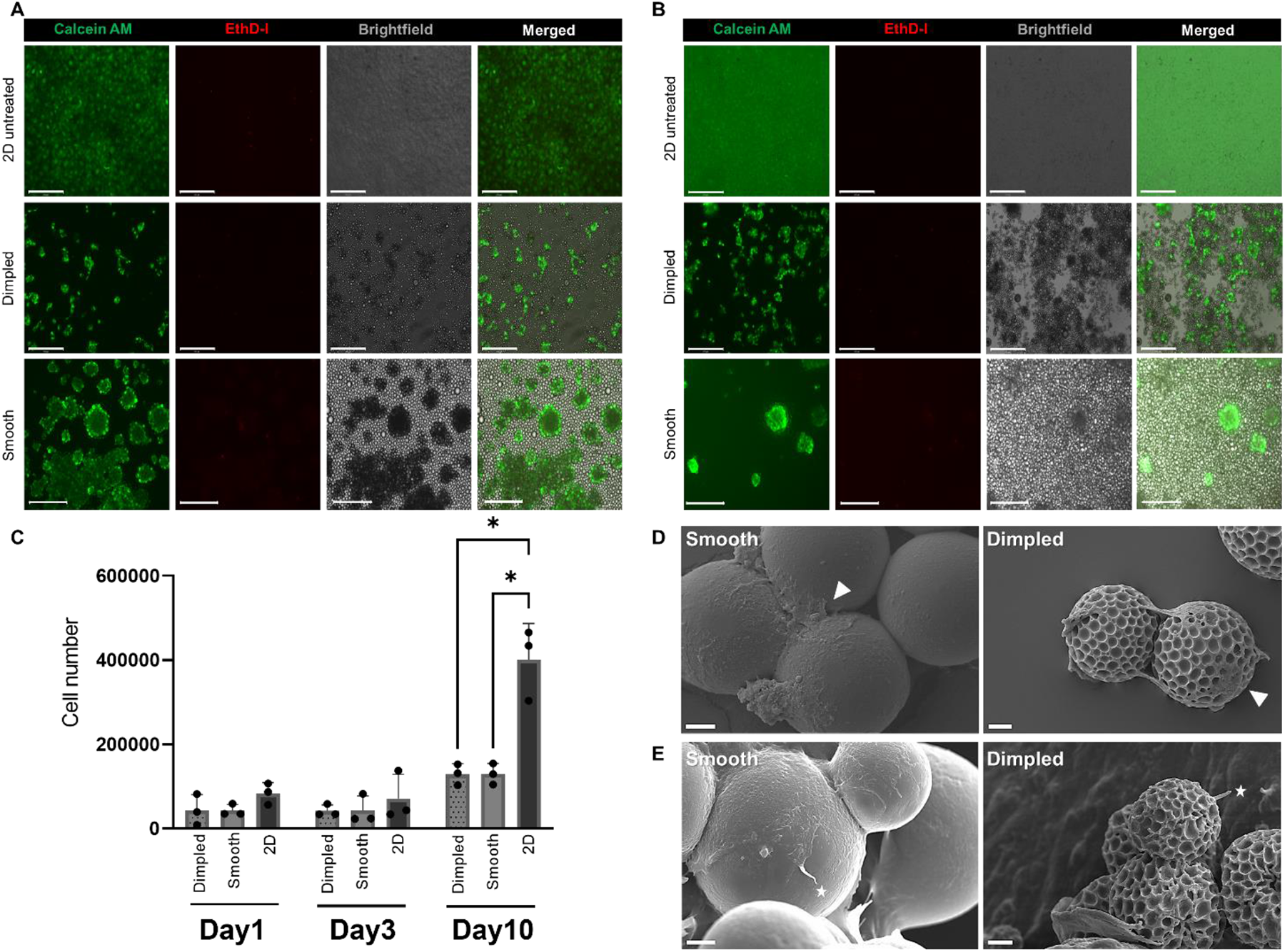
Viability, proliferation, and morphology of C3H10T1/2 cells on smooth versus dimpled microparticles. (A, B) Representative fluorescence microscopy images showing high viability of C3H10T1/2 cells on smooth and dimpled microparticles 3 days after seeding in (A) serum-reduced (2% FBS) culture media and (B) in serum-containing (10% FBS) media. Live cells are stained with calcein-AM (green) and dead cells with ethidium homodimer III (EthD) (red).) (Scale bar=: 200 μm). (C) Proliferation of C3H10T1/2 cells assessed using PrestoBlue on dimpled and smooth microparticle at days 1, 3 and 10 days after seeding. Statistical significance calculated with two-way ANOVA with Tukey’s multiple comparisons test. Data represents mean ± standard deviation for three independent repeats. (*p < 0.05). (D, E) Scanning electron microscopy images showing the morphologies of C3H10T1/2 cells 3 days after seeding on the smooth and dimpled microparticles. White arrowheads in (D) indicate C3H10T1/2 cells attached to microparticles. White stars in (E) indicate representative cytoplasmic protrusions observed on cells cultured on microparticles (Scale bar= 10 μm). Abbreviations: Calcein-AM, calcein-acetoxymethyl; EthD-I, ethidium homodimer III

Cell proliferation was evaluated at days 1, 3, and 10 using the PrestoBlue assay (Figure 2C). Whilst cell numbers increased on dimpled and smooth microparticles over 10 days of culture, there were no significant differences observed between cell numbers on the two microparticle designs across all time points. However, cell numbers in 3D-cultured samples were significantly lower than 2D-cultured controls at day 10 (*p* ≤0.05).

Cells displayed different morphologies when cultured on different topographical designs (Figure 2D). Cells adopted more elongated morphologies on dimpled microparticles, whilst cells were more spread out on the smooth surfaces. Interestingly, we observed protrusions projecting from the surface of cells cultured on the microparticles (Figure 2E and Figure S2). On smooth microparticles, these slender cellular protrusions displayed an average length of 11.21 ± 2.11 µm (*n*=3), whereas on dimpled microparticles these protrusions were noticeably thicker and shorter, with an average length of 5.59 ± 0.83 µm (*n*=6).

### Dimpled topographical features alter focal adhesion assembly

The actin cytoskeleton is a dynamic structure that determines cellular shape and regulates its behaviour, with focal adhesions being the primary sites of cell adhesion to a substrate. The formation of stable links with the extracellular matrix leads to integrin clustering and organisation into focal adhesions [33]. To investigate how the arrangement of cytoskeletal proteins was influenced by the changes in 3D topographical features, the organisation of F-actin and vinculin (a membrane cytoskeletal protein found in focal adhesions) in response to different microparticle designs was examined after 3 days of culture and compared to 2D-cultured controls (Figure 3).

**Figure 3:**
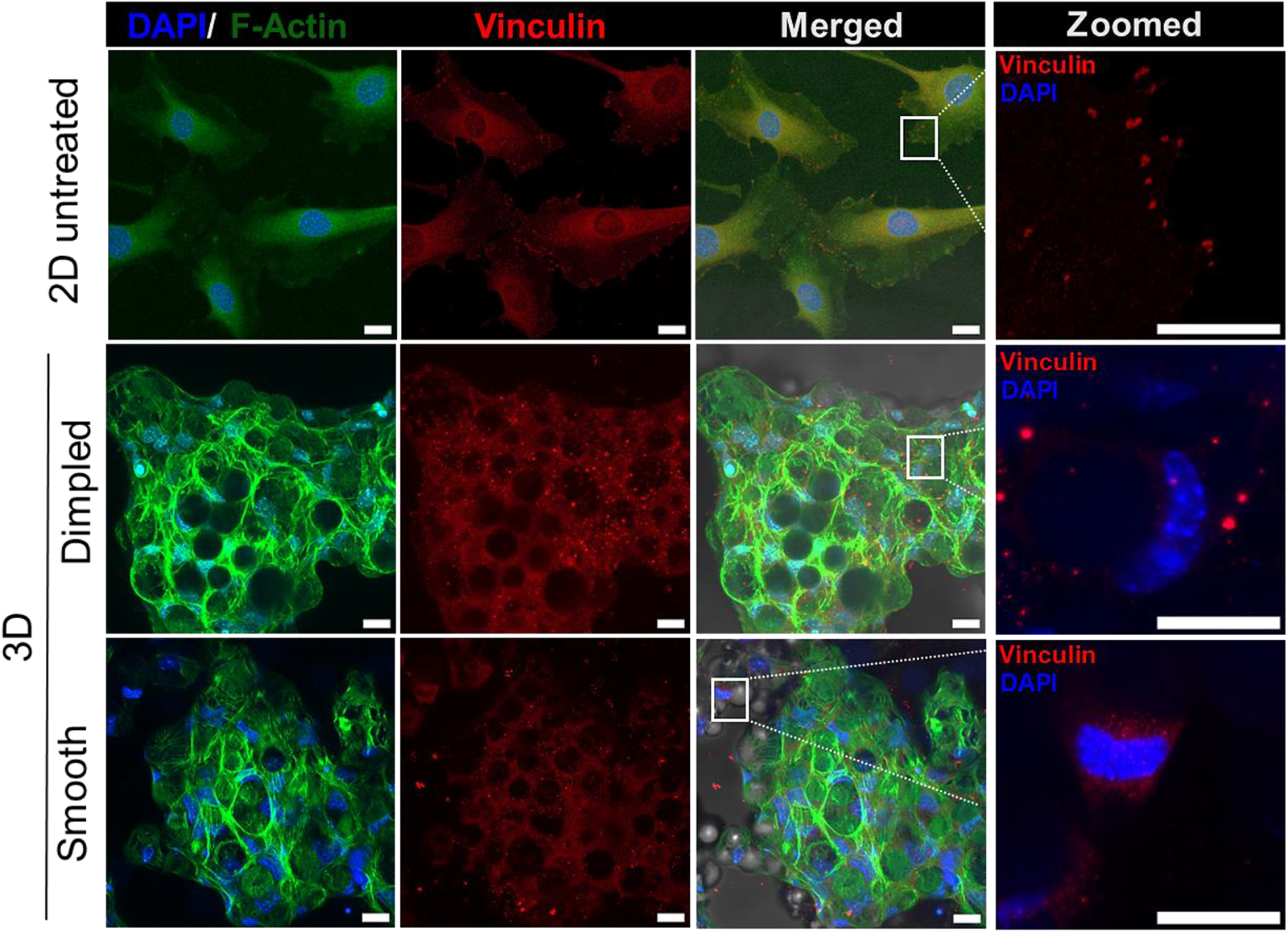
Three-dimensional topographical features display a differential effect on focal adhesion organisation. Representative immunofluorescence images of C3H10T1/2 cells showing differences in cytoskeletal organisation, stained for vinculin (focal adhesion marker; red), F-actin (green) and nuclei (DAPI; blue). The rightmost panel shows an enlarged view of the vinculin staining observed in the square inset in the merged images. Cells were cultured on two microparticle designs and 2D controls in serum-reduced culture media for 3 days (Scale bar = 20 μm). Abbreviations: DAPI, 4’,6-diamidino-2-phenylindole.

Cell culture on microparticles demonstrated the formation of focal adhesions linked to actin stress fibres (Figure 3). On the 2D-cultured planar control, cells cultured in serum-reduced medium displayed well-defined focal adhesions, with distinct vinculin-positive streaks organised at the leading edge of the cell but also dispersed widely in the cytoplasm. Network-like actin microfilaments with large stress fibres were observed in 3D-cultured cells (Figure 3). Perinuclear vinculin-containing focal adhesion sites were visible in cells cultured on smooth microparticles (Figure 3). In contrast, vinculin was not organised into focal adhesions in cells cultured on dimpled microparticles, but was found distributed throughout the cytoplasm, which may suggest an initial state of F-actin polymerisation [34] or depolymerisation of the actin filaments [35]. This demonstrated that the changes in focal adhesion assembly were correlated with the changes observed in cell morphology.

### Dimpled topographical features induce primary cilia formation

As primary cilium formation has been linked with cellular morphology and organisation of cytoskeletal network [36], 2D and 3D microparticles-based cultures were stained for the ciliary membrane protein ARL13B, a small regulatory Ras GTPase highly enriched in primary cilia [37]. Cilia formation can be affected by sub-micron topographies (i.e. <10 µm) in hMSCs [38], and we therefore investigated cilia formation in C3H10T1/2 cells cultured on microparticles using ARL13B immunostaining.

Cells cultured on dimpled microparticles displayed ARL13B^+^ primary cilia, which were absent in cells cultured on smooth microparticles and few in number in 2D-cultured untreated controls (Figure 4). To the best of our knowledge, this is the first report of the presence of primary cilia in 3D microparticle-based cultures.

**Figure 4:**
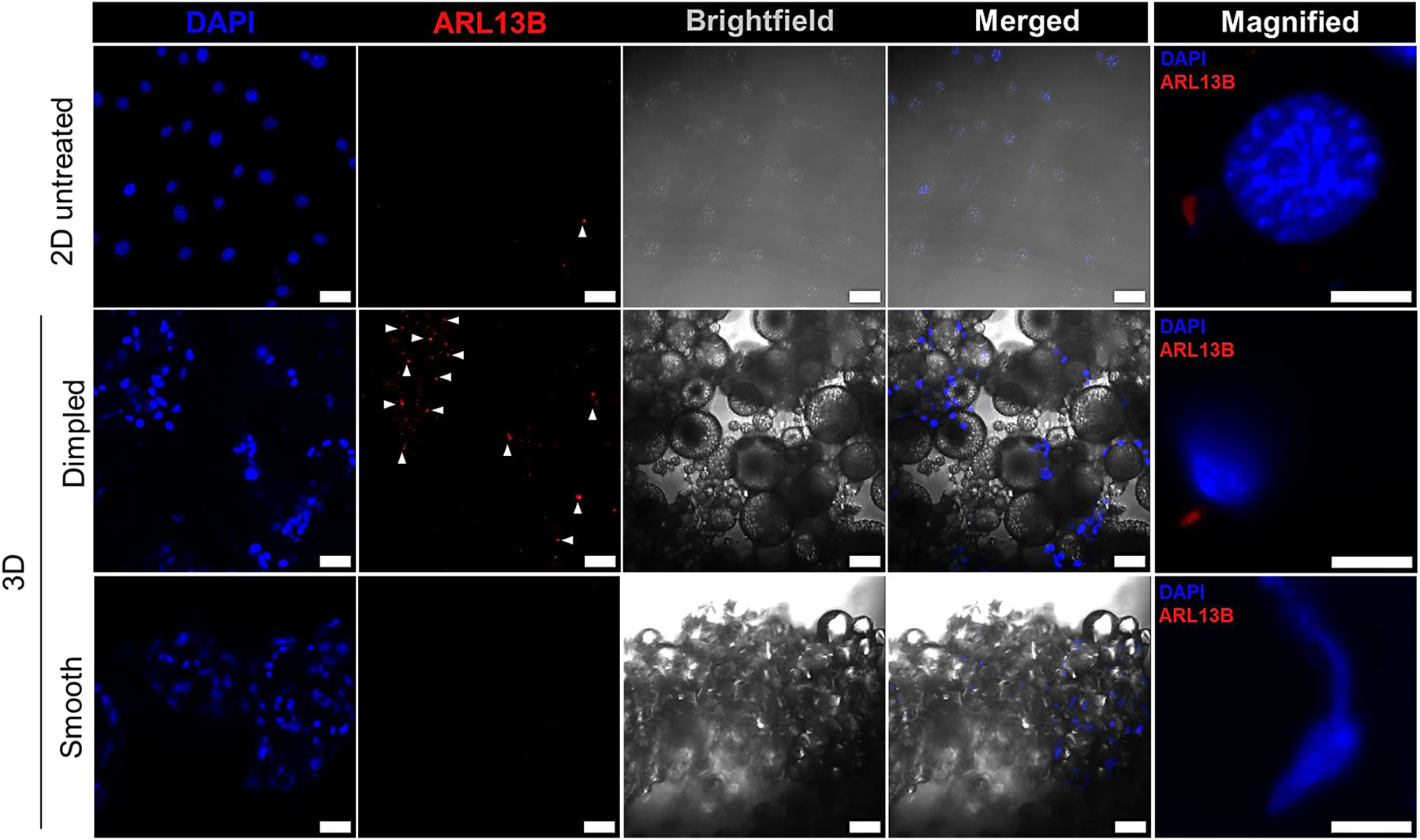
Representative maximum intensity projection immunofluorescence images of C3H10T1/2 cells stained for ARL13B (ciliary marker; red) and nuclei (DAPI, blue), with positive ciliary staining indicated by white arrows. Cells were cultured on microparticles and 2D untreated cultures in serum-reduced media for 3 days. (Scale bar = 50μm). Higher magnification images of ciliary staining are also included as “Magnified” (Scale bar = 10μm). Abbreviations: DAPI, 4’,6-diamidino-2-phenylindole; ARL13B, ADP-ribosylation factor-like protein.

### Dimpled topographical features induce osteogenic differentiation of C3H10T1/2 cells without the addition of exogenous osteoinductive factors

Given the presence of primary cilia when culturing C3H10T1/2 cells on dimpled microparticles, we next investigated the effects of dimpled topographical features on the primary cilium-associated Hedgehog (Hh) signalling pathway, and how this may influence downstream osteogenic differentiation on these osteo-inductive topographical features [2]. To confirm the osteo-inductive capability of dimpled microparticles using the C3H10T1/2 mesenchymal progenitor cells, we quantified early and late markers of osteogenesis (*Runx2* and *Bglap2* encoding osteocalcin, respectively; Figure 5A-D). Since Runx2 binds to promoter sequences of bone-specific proteins to regulate osteogenesis [39], Runx2 activation indicates early osteogenic commitment. Positive control cultures were treated with either 2µM purmorphamine or osteo-inductive media, chemical inducers of *Runx2* [40] and *Bglap2* [41] expression, respectively (Figure S3).

**Figure 5:**
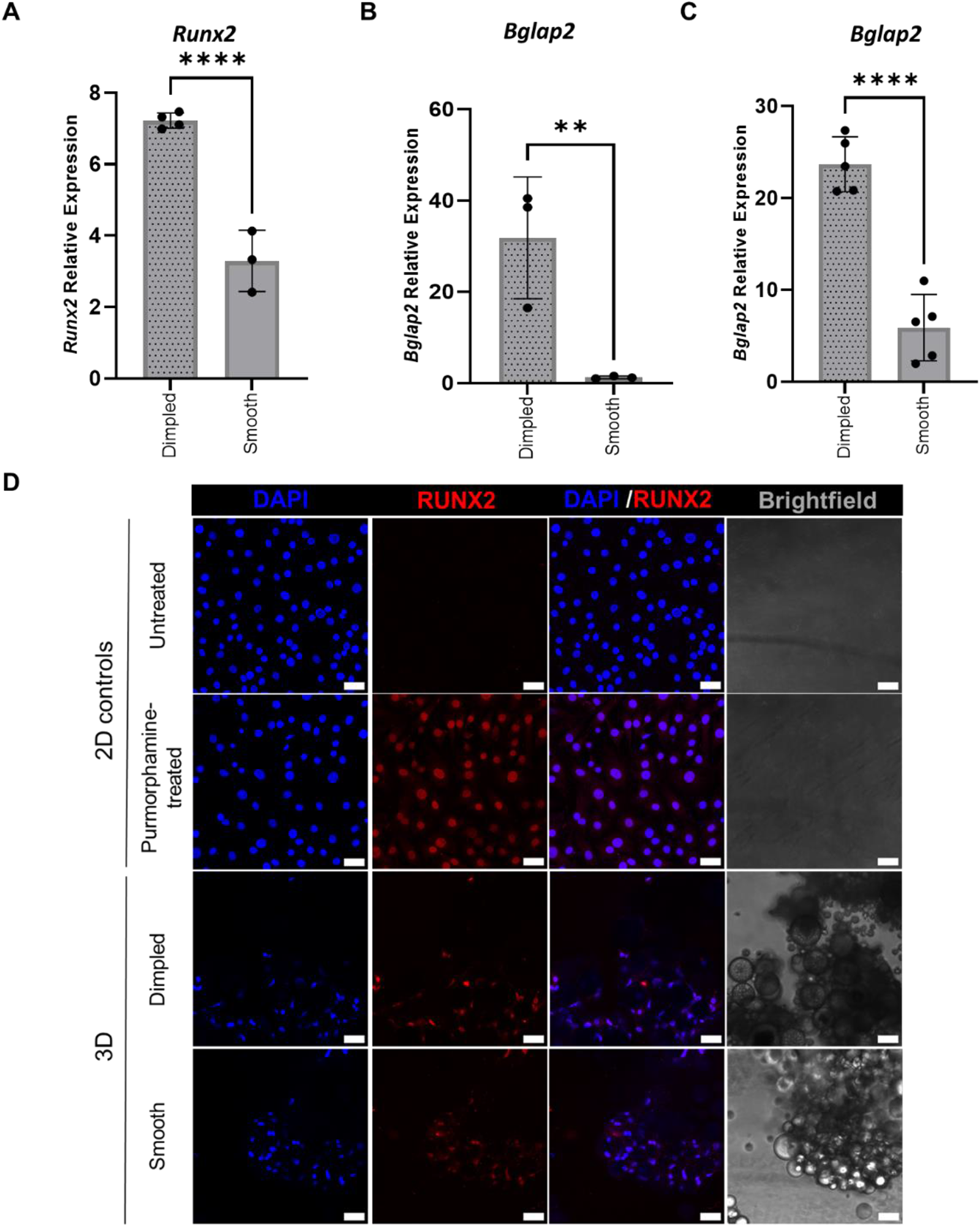
Culture on topographically-textured microparticles induces the expression of early and late markers of osteogenesis without the addition of exogenous osteoinductive factors. (A-C) Relative quantitative real-time PCR (qPCR) analysis of Runx2 expression levels after 3 days (A) and Bglap2 expression levels after 3 days (B) and (C) 7 days in in serum-reduced media, represented as fold change normalised to Gapdh and relative to 2D untreated controls. Statistical significance calculated with one-way ANOVA with Tukey’s multiple comparisons test. Values are means ± standard deviation for a minimum of 3 independent biological repeats. (*p < 0.05, **p < 0.01, ****p< 0.0001). (D) Representative maximum projection images obtained by confocal microscopy of C3H10T1/2 cells stained for RUNX2 (red) after 3 days (Scale bar = 50 μm) and nuclei (DAPI; blue). 2D-cultured positive controls were treated with 2µM purmorphamine for 3 days to induce Runx2 expression. Abbreviations: DAPI, 4’,6-diamidino-2-phenylindole; Runx2, Runt-related transcription factor2; Bglap2, bone gamma-carboxyglutamate protein 2; Gapdh, Glyceraldehyde-3-Phosphate Dehydrogenase

Cells cultured on dimpled microparticles displayed a statistically significant increase in *Runx2* expression levels compared to those cultured on smooth microparticles (7.26-fold versus 3.91-fold; *p* <0.0001) relative to 2D negative controls and normalised to *Gapdh* (Figure 5A). To further confirm the osteoinductive effect of 3D dimpled topographies on C3H10T1/2 cells without the use of exogenous biochemical supplements, relative expression analysis of *Bglap2* (a mature osteoblast marker) was determined 3 and 7 days after seeding (Figure 5B and 5C). Culture on dimpled microparticles significantly upregulated the expression levels of *Bglap2* by 31.87-fold (*p* <0.01) in C3H10T1/2 cells after 3 days in culture, whereas no significant differences in the levels of *Bglap2* expression by cells cultured on smooth microparticles was observed relative to 2D untreated controls (Figure 5B). This significant increase in *Bglap2* expression on dimpled microparticles continued to be observed at day 7, with a 23.39-fold increase of *Bglap2* expression levels relative to 2D negative controls. Cells cultured on smooth microparticles for 7 days displayed 8.21-fold increase in *Bglap2* expression relative to 2D untreated controls (*p* <0.0001) (Figure 5C).

Immunostaining was also carried out at day 3 to confirm the effect of dimpled topography on RUNX2 expression. Immunostaining also confirmed notably higher expression levels of RUNX2 in C3H10T1/2 cells cultured on dimpled microparticles relative to smooth microparticles. Nuclear localisation of RUNX2 was observed on dimpled microparticles, whereas RUNX2 was retained mostly in the cytoplasm in cells cultured on smooth microparticles (Figure 5D). These findings confirmed the enhanced ability of dimpled microparticles to induce osteogenic differentiation of C3H10T1/2 cells without employing exogenous osteo-inductive factors.

### Topographically-induced osteogenesis is associated with the Hedgehog signalling pathway

The Hedgehog signalling pathway is essential in stem cell regulation and development [42]. Given the well-known role of Hh signalling pathway in driving MSCs osteogenic commitment by increasing the expression levels of *Runx2* [43] and boosting osteoblast maturation by upregulation of the expression of osteocalcin [44], we next investigated the involvement of the Hh signalling pathway in the 3D dimpled topography-induced osteogenesis. The expression of Hh signalling pathway-related genes were quantified at day 3 after seeding (Figure 6).

Interestingly, the trends seen in canonical Hh induced gene expression levels in cells cultured on dimpled microparticles were similar to those observed for osteogenic gene expression at day 3 post-seeding. Expression levels of *Gli1* and *Ptch1,* hallmark target genes of canonical Hh signalling, were significantly upregulated on dimpled microparticles by 17.31-fold (p <0.0001, Figure 6A) and 2.06-fold (p <0.01, Figure 6B), respectively, relative to 2D untreated cultures. Remarkably, dimpled microparticles were able to induce *Gli1* expression to a comparable level to that observed in 2D positive controls treated with 2µM purmorphamine, a *Smo* agonist, which showed a 21.54-fold increase in *Gli1* expression relative to the corresponding 2D-vehicle only controls (Figure 6A). Moreover, the expression level of Smo, a central transducer of the Hh signal that is not a target of Gli-dependent transcription [45], was also increased by 12.99-fold in the cells cultured on the dimpled microparticles (*p* <0.0001, Figure 6C). Expression levels of the three Hh signalling-related genes showed no statistically significant differences between smooth microparticles and 2D negative controls. GLI1 protein expression was verified by western blotting and quantified by densitometry analysis at day 3, showing a 25.90-fold increase in GLI1 expression in cells cultured on dimpled microparticles relative to its expression in 2D untreated controls (Figure 6D).

High GLI1 protein expression was observed on the C3H10T1/2 cells cultured on dimpled microparticles by immunofluorescence staining, similar to that seen in the purmorphamine-treated cells in 2D cultures (Figure 6E). In contrast, no expression of GLI1 was observed in C3H101/2 cells when cultured on smooth microparticles. This was consistent with the results observed using RT-qPCR and western blotting.

Taken together, this data demonstrates that dimpled topographically-textured microparticles induced the activation of Hh signalling and osteogenesis in the mesenchymal progenitor C3H10T1/2 cells without the addition of exogenous biochemical additives or Hh pathway agonists.

**Figure 6:**
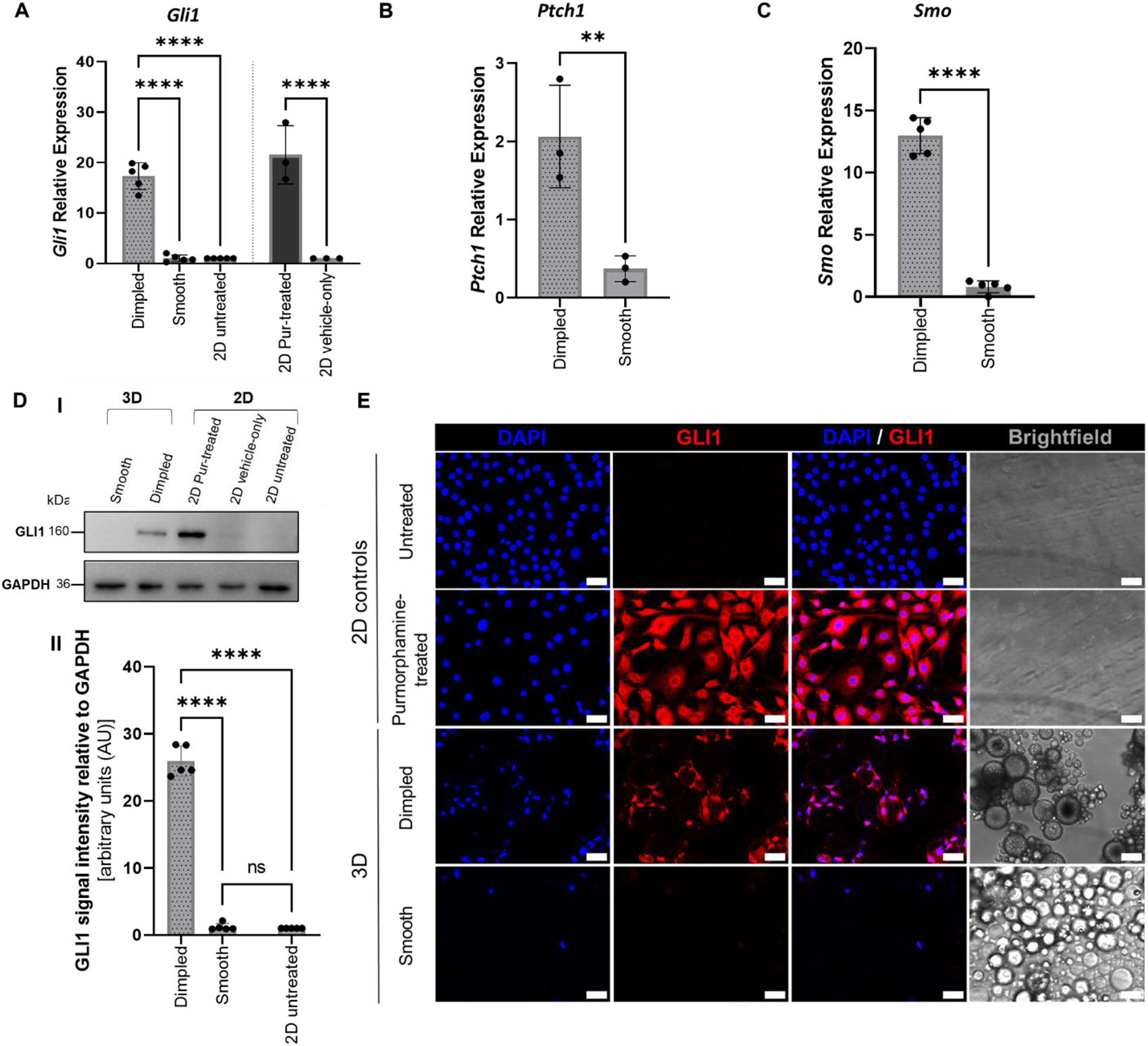
Dimpled microparticles upregulate the expression levels of Hh-related markers. Relative quantitative real-time PCR (qPCR) analysis of Hh-related genes (A) Gli1, (B) Ptch1 and (C) Smo after 3 days in culture in serum-reduced media, represented as fold change normalised to GAPDH and relative to 2D controls. Expression levels in 2D purmorphamine-treated controls (2µM) were calculated relative to 2D-vehicle only controls (treated with 0.06% DMSO; n=3). (D) Representative Western blot (I) and densitometry analysis performed (II) showing GLI1 expression levels in C3H10T1/2 cells 3 days post-seeding, normalised to GAPDH (n=5). Statistical significance calculated with one-way ANOVA with Tukey’s multiple comparisons test for all data, with the exception of 2D controls where unpaired Student’s t-test was used. Values are shown as means ± standard deviation (*p < 0.05, **p < 0.01, ***p< 0.001, ****p< 0.0001). (E) Representative confocal immunofluorescence microscopy images of C3H10T1/2 cells immunostained for GLI1 (red) and nuclei (DAPI; blue). 2D positive controls were treated with 2µM purmorphamine (Scale bar = 10μm). Abbreviations: DAPI, 4’,6-diamidino-2-phenylindole; Gli1, glioma associated oncogene homolog 1; Ptch1, Patched1; Pur-treated, purmorphamine-treated; Smo, Smoothened; Gapdh, Glyceraldehyde-3-Phosphate Dehydrogenase.

### Osteogenesis Induced by Dimpled Topography is Mediated by Smo-Dependent Hh Signalling Pathway

Activation of canonical Hh signalling requires regulated trafficking of proteins into and out of the primary cilium [46]. During pathway activation, Smo accumulates in primary cilia to signal activation of Gli2 and Gli3, which then induce the expression of Gli-target genes, among which Gli1 and Ptch1 are widely recognised [47]. Therefore, we further investigated the role of Smo in the topographically-induced activation of the Hh signalling pathway and osteogenesis. We hypothesised that dimpled topographies sensed by C3H10T1/2 cells lead to Smo-dependent activation of Hh signalling, which upregulates downstream osteogenesis-related genes expression. For this purpose, we quantified the expression levels of *Gli1* and osteogenesis-related genes, *Runx2* and *Bglap2*, with and without the Smo inverse agonist, KAAD-cyclopamine (Figure 7). The concentration in DMSO was optimised to preserve the textured topographical features of the fabricated microparticles (Figure S4).

**Figure 7:**
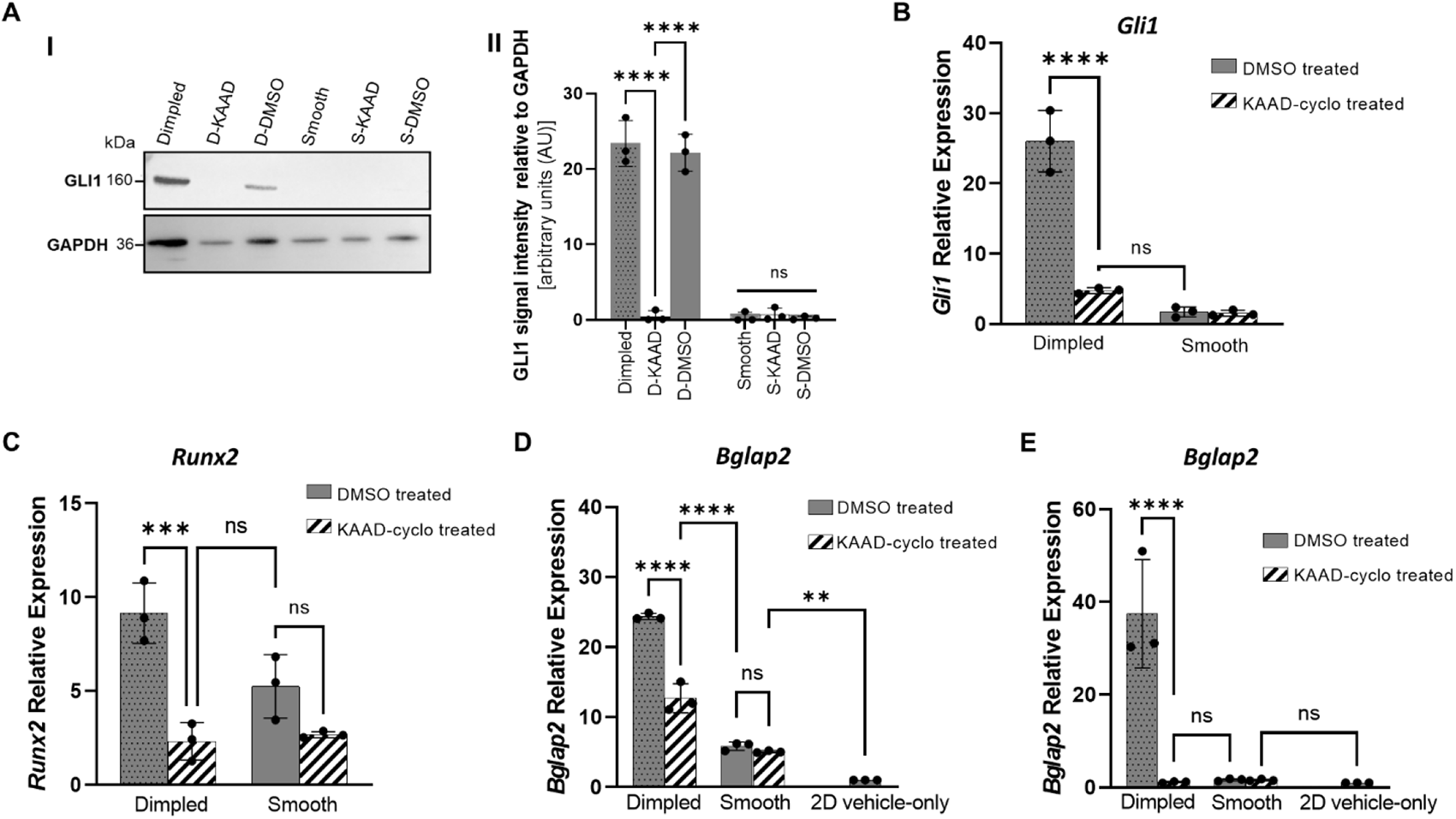
Hedgehog activation and topographically induced osteogenesis is Smo-dependant. (A) Representative Western blot (I) and densitometry analysis (II) showing GLI1 expression levels in C3H10T1/2 cells after 3 days of 300µM KAAD-cyclopamine treatment, normalised to GAPDH (n=3). (B-D) Relative qPCR analysis of (B) Gli1, (C) Runx2 and (D, E) Bglap2 after 3 (D) and 7 (E) days in culture, represented as fold change normalised to GAPDH and relative to 2D vehicle-only controls. Statistical significance was determined using one-way ANOVA with Tukey’s multiple comparisons test. Values are means ± standard deviation for three independent biological repeats. (**p < 0.01, ***p< 0.001, ****p< 0.0001, ns= no statistically significant difference). Abbreviations: D-KAAD and S-KAAD, 300µM KAAD-cyclopamine treated cells cultured on dimpled and smooth microparticles, respectively: D-DMSO and S-DMSO, 0.06% DMSO-treated cells cultured on dimpled and smooth microparticles, respectively; Gli1, glioma associated oncogene homolog 1; Runx2, Runt-related transcription factor2; Bglap2, bone gamma-carboxyglutamate protein 2; DMSO, dimethyl sulphoxide; KAAD-cyclo, KAAD-cyclopamine; GAPDH, Glyceraldehyde-3-Phosphate Dehydrogenase

KAAD-cyclopamine treatment resulted in significant decrease in the expression of all three genes under investigation in the cells cultured on dimpled microparticles. Inhibition of Smo using KAAD-cyclopamine significantly prevented *Gli1* upregulation in cells cultured with dimpled topographies at both the transcript and protein levels (*p* < 0.0001, Figure. 7A and 7B). In the presence of KAAD-cyclopamine, no statistically significant differences in the expression levels of *Gli1* in C3H10T1/2 cells cultured on smooth microparticles were observed.

Pharmacological inhibition of Smo also prevented the previously observed upregulation of osteogenesis-related genes in C3H10T1/2 cells cultured on dimpled microparticles. This was demonstrated by the statistically significant reduction in expression levels of *Runx2* (*p* <0.001, Figure 7C) and *Bglap2* in KAAD-cyclopamine treated cells (*p* <0.0001, Figure 7D and E). The expression levels of *Bglap2* at day 7 in the presence of KAAD-cyclopamine treatment were significantly reduced to the levels observed in C3H10T1/2 cells cultured on smooth microparticles (*p* <0.0001, Figure 7E). Taken together, our results show that the topographical design of microparticles can differentially activate the canonical Hh signalling pathway. Furthermore, our data demonstrates the modulation of osteoinduction, mediated by the canonical Smo-dependent Hh signalling pathway, through engineering cell-instructive 3D microenvironments on topographically-textured microparticles.

## Discussion

Modularity is ubiquitous in biological systems. Polymeric microparticles offer great potential as functional biomaterials-based building blocks for use in bottom-up tissue engineering and advanced culture systems. Topographically-textured microparticles are a powerful tool to direct cell response, and can be customised to add further functionality [48], enhancing their utility. Providing mechanistic insights into how topographical design criteria for microparticles-based environments influence cell signalling opens the door for *ex vivo* control of cell fate in disease model development. Therefore, the aim of this study was to investigate the Hh signalling-mediated osteogenic differentiation of 3D micro-topographical cues using topographically-textured PLA microparticles in the absence of exogenous agonists. We demonstrate that the osteo-inductive influence of C3H10T1/2 mesenchymal progenitor cells by dimpled microparticles is mediated by the activation of canonical Hh signalling, as demonstrated by the upregulation of various Hh-related signalling markers and increased primary ciliogenesis, demonstrating the Smo-dependency of osteogenic marker expression (Figure 8).

When employing microparticles as culture substrates, cell viability and proliferation are critical. PLA-based microparticles were employed in this study to avoid the degradation and loss of the engineered topographical features in culture [2]. C3H10T1/2 cells showed excellent attachment and viability on the microparticles. Additionally, increased proliferation rates in 2D controls compared to their 3D counterparts was observed. This aligns with previous studies reporting that cells display reduced proliferation rates in 3D cultures compared to those cultured in 2D, and are matrix-dependent [2,49–51]. Proliferation rates in 3D-cultured systems better represent in vivo models than 2D cultures [52]. The differences in C3H10T1/2 cell morphologies when cultured on smooth and dimpled microparticles agree with those we previously reported using human MSCs [2], which are hypothesised to be due to preferential use of different mechano-transducers to adapt to the different physical microenvironments [53]. Human MSCs attachment on dimpled microparticles was reported to be mediated primarily via integrins α_5_ and α_v_β_3_, unlike those cultured on smooth microparticles [2]. Integrins are known to connect to the primary cilium via the actin cytoskeleton, and focal adhesions are clusters of integrins and transmembrane receptors that mechanically link the ECM to the actin cytoskeleton [54,55]. Remarkably, C3H10T1/2 cells also displayed cytoplasmic projections extending from the surface of cells cultured on dimpled microparticles (Figure 2E). These may be similar to the ‘micro-spikes’ observed with synovial mesenchymal stem cells reported in a recent study, which were suggested to have roles in mechanosensing and cell-matrix interactions [56].

Changes in topography are known to induce actin cytoskeletal changes [57]. Vinculin is a ubiquitously expressed mechanosensitive protein that interacts with F-actin during the recruitment of actin filaments to growing focal adhesions, as well as being involved in capping of actin filaments to regulate actin dynamics [58]. We demonstrated that C3H10T1/2 cells cultured on smooth surfaces showed diffuse perinuclear vinculin staining (Figure 3), in agreement with previous studies on C3H10T1/2 and human fibroblasts h-TERT immortalised BJ1 cells [59,60]. Zhoa *et al.* reported that C3H10T1/2 cells displayed reduced levels of vinculin staining on hydroxyapatite-based multilayers, resulting in enhanced osteogenic differentiation [59]. The disorganised focal adhesion staining observed in C3H10T1/2 cells cultured on dimpled topographies is visibly similar to the disruption of focal adhesion assembly observed with oestrogen withdrawal in murine osteocyte-like cells, which resulted in primary cilia formation [54]. Micro-scale topographical features can block cell spreading by directly interfering with the establishment and maturation of focal adhesions [61]. Disrupted focal adhesions leads to actin depolymerisation and inhibit actin contractility [54,62], which was found to promote ciliogenesis [63]. Furthermore, Zhang *et al.* reported that introducing microscale topographies to 2D discs induced osteogenesis of hMSCs and caused changes in actin organisation coupled with ciliogenesis [38]. Therefore, the 3D dimpled topographies may cause differential assembly of focal adhesions in smooth versus dimpled microparticles, leading to cytoskeletal remodelling and ciliogenesis. This aligns with previous research reporting that actin cytoskeleton organisation can be modulated by designing specific surface topographical features that mimic ECM properties [57,64].

The formation of primary cilia exclusively in C3H10T1/2 cells cultured on dimpled microparticles was confirmed by immunostaining for ARL13B, a ciliary membrane marker (Figure 4). Ciliogenesis can be promoted via changes in expression of actin regulatory factors required for branched actin network formation [65]. Changes in cell morphology can also have a strong influence on the formation of cilia [66,67]. It has been reported that cilia formation and length can be affected by sub-micron topographies (i.e. less than 10µm) in MSCs [38]. The primary cilium is known to be essential for the transduction of the Hh signalling pathway in vertebrates [46,68], with important roles in mesenchymal cell mechanobiology and mechanotransduction [35,69,70]. Alterations in primary cilia structure or function directly influences osteogenesis via the modulation of Hh signalling pathway [54], which is indispensable in the early stages of osteogenesis and subsequent osteoblast maturation [12,71].

Our data demonstrated that C3H10T1/2 mesenchymal progenitor cells cultured on dimpled microparticles underwent osteogenesis in the absence of any exogenous supplements, which was confirmed by the increase in the expression levels of osteogenesis-related genes, including *Runx2* and *Bglap2* compared to cells cultured on smooth microparticles and 2D controls (Figure 5). Runx2 is an important marker of early osteogenic differentiation [19,72]. Activation of Hh signalling pathway leads to direct induction of *Runx2* transcription [73], which further activates the Hh pathway through the induction of *Gli1* and Ptch1 [74]. Expression of Bglap2, which serves as a late marker of osteogenic differentiation [75], was also upregulated. Culture of C3H10T1/2 cells on dimpled microparticles displayed a significant increase in *Bglap2* expression at day 3 post-seeding, which persisted to day 7. This aligns with our previous study using human MSCs, where we demonstrated that topographically-textured 3D microparticles can induce osteogenic differentiation of human MSCs *in vitro* [2]. Though PLA lacks intrinsic osteo-inductivity [76,77], previous studies have reported that stiffness can stimulate osteogenic differentiation to some degree [2,78]. This can explain the increase in expression levels of *Bglap2* also observed on smooth microparticles at day 7 post-seeding (Figure 5). However, this was still significantly lower than expression levels of these markers by the cells cultured on dimpled microparticles.

Increasing evidence has demonstrated the crucial roles of Hh signalling in various bone development processes [79–81]. It is well-known that Hh signalling is mechano-responsive and required for loading-induced osteogenesis in MSCs [12]. Recent studies have increasingly focused on the potential role of the Hh pathway in response to mechanical cues [12,82–86]. The influence of different mechanical cues on Hh signalling pathway has also been reported, such as cyclic tensile strain [82–84], hydrostatic loading [85,86], oscillatory fluid flow [12] and matrix stiffness [87]. Previous studies that have reported the influence of topographical features on Hh signalling activation were mostly conducted using planar titanium surfaces [19,88]. Recent studies have shown that introducing planar topographies on titanium surfaces increased the expression of Hh-related genes [19,88,89]. Our data showed that dimpled microparticles significantly upregulated the activity of the canonical Hh pathway, demonstrated by the increase in expression levels of *Gli1* and *Ptch1,* and supported by the upregulation of *Smo* levels compared to smooth microparticles and 2D cultures (Figure 6). This causal relationship between Hh pathway activation and osteogenic gene expression demonstrated here is consistent with previous findings, which showed increased expression of *Runx2* and *Bglap2* when the Hh pathway was chemically activated in MSC by purmorphamine treatment [40,90] and by using a recombinant N-SHH, which increased ALP activity^91^. Remarkably, dimpled microparticles enhanced Hh signalling at day 3 after seeding to a level similar to that observed for 2D cultures treated with 2µM purmorphamine (Figure 6). This highlights the potential of employing dimpled microparticles as a physical alternative to biochemical Hh agonists in discovery research. The increased expression of Smo observed in dimpled microparticles agrees with a previous study that utilised rough titanium surfaces to culture osteoblast-like MG63 cells [19], reporting that surface topographies enhanced the expression levels of Hh-related genes including *Smo*. This represents an exciting avenue for future research to shed further light on the mechanism by which topography enhances Smo expression. It is well-established that Smo in the presence of primary cilia can drive activation of Hh signalling [47]. Smo translocates to the primary cilia in response to Hh ligands, with its activation promoting the release of Gli proteins from their carrier protein, Suppressor of Fused (Sufu). Direct transcriptional targets of Hh signalling include Gli1 and Ptch1 [53,92] (Figure 8).

To demonstrate the role of Hh signalling in the osteo-inductive capabilities of 3D dimpled topographies investigated herein, we suppressed canonical Hh signalling using the Smo inhibitor KAAD-cyclopamine, subsequently quantifying osteogenesis-related gene expression. Our results demonstrated that KAAD-cyclopamine suppressed Hh activity and prevented the expression of the osteogenesis-related genes (*Runx2* and *Bglap2*) in C3H10T1/2 cells cultured on dimpled microparticles (Figure 7). This data confirmed our hypothesis that Hedgehog–Gli signalling mediates the osteo-inductive impact of dimpled microparticles on C3H10T1/2 cells. Unlike dimpled microparticles-based cultures, the expression levels of *Bglap2* observed at day 7 with smooth microparticle cultures is independent of Smo. This suggests the involvement of different osteogenesis-related signalling pathways in the promotion of osteogenesis, such as Wnt, TGFβ, bone morphogenic protein (BMP) or Notch signalling pathways, potentially mediated by PLA surface stiffness [93–95].

In summary, we provide novel mechanistic insights into the osteo-inductive influence of dimpled topographical design of PLA microparticles, demonstrating that the osteoinductive effect of these dimpled microparticles is mediated by Hedgehog signalling, and that the route of activation of the Hh-Gli1 pathway is Smo-dependent (Figure 7). These micron-sized, readily injectable surface-engineered microparticles offer significant potential in bone regenerative applications and can be used in modelling diseases for which Hh signalling dysfunction is implicated without the need for adding exogenous agonists.

**Figure 8:**
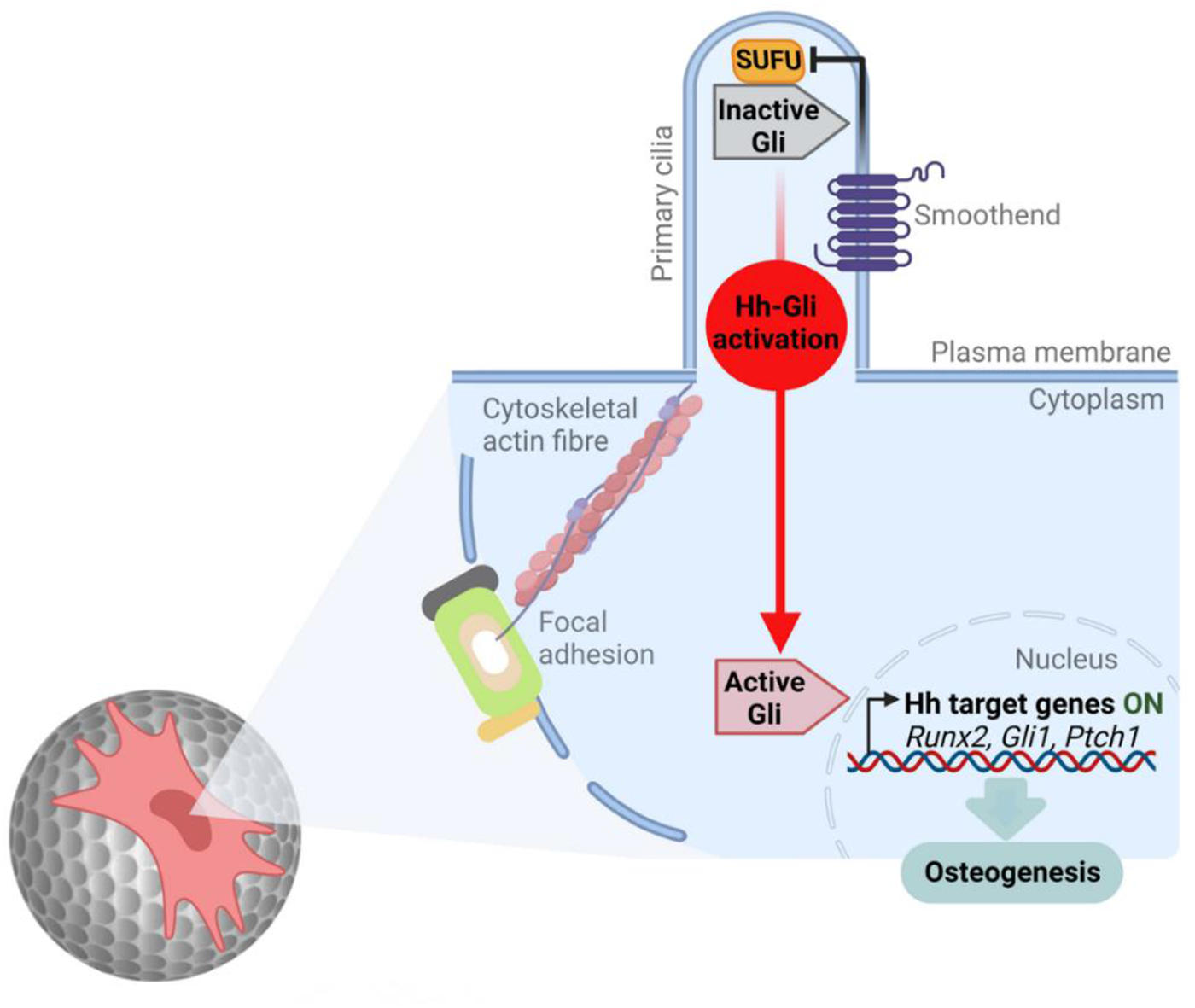
Schematic depicting the proposed mechanism underpinning the osteoinductive influence of 3D dimpled surface topographical features of PLA microparticles on C3H10T1/2 cells. Dimpled topographies are sensed by C3H10T1/2 cells, which leads to changes in cytoskeleton organisation and the formation of primary cilia. This facilitates the activation of the Hedgehog signalling pathway and translocation of active Gli1 into the nucleus, where it upregulates the expression of Hh-and osteogenic-related genes, leading to the expression of early and late markers of osteogenesis. Image was created using Biorender.com. Abbreviations: Hh, Hedgehog signalling pathway; Gli1, glioma associated oncogene homolog 1; Runx2, Runt-related transcription factor2; Ptch1, Patched 1

## Conclusion

This study uncovers novel mechanistic insights into the osteo-inductive impact of 3D topographical patterning of microparticles on the modulation of Hh signalling – a key pathway in cellular differentiation. Our findings will provide key guiding parameters for designing microparticle-based models that mimic key cellular and molecular regulation of Hh signalling in bone development and regeneration.

There is growing interest in the development of advanced culture substrates that can direct endogenous signalling pathways. Currently, researchers need to compromise between complexity and control in *in vitro* stem cell differentiation. When control over differentiation is desired, only limited complexity in cellular composition and culture media supplementation is possible. Alternatively, a high degree of complexity in cellular composition limits the ability to control individual cell fates, typically due to the need for utilisation of a common culture media. Tailoring the architectural features of microparticles as modular culture substrates will allow the assembly of cell-microparticle aggregates as modular building blocks. Modular scaffolds can be prepared by assembling varying topographical designs of polylactic acid-based microparticles as building blocks to create novel Hh-related disease model systems free from confounding biochemical factors that are typically added to induce differentiation *in vitro*. This will open avenues for engineering spatially controlled, heterogeneous tissue constructs as advanced *in vitro* experimental systems suitable for *in vitro* discovery research and high-throughput screening of novel drug candidates for treating diseases for which Hh signalling dysfunction is associated.

## Acknowledgements

MA acknowledges funding by a University Academic Fellowship (University of Leeds, UK). FG is supported by a scholarship from Kuwait University, Kuwait. We thank Dr Ruth Hughes from the Faculty of Biological Sciences Bioimaging facility for support with the Zeiss LSM880 Airyscan confocal microscope, funded by a Wellcome Trust (WT104918MA). We thank John Harrington and Stuart Micklethwaite from Leeds Electron Microscopy and Spectroscopy Centre (LEMAS) for their support in this work. Finally, we acknowledge Tayah Hopes for her invaluable support with RT-qPCR, Dr Iain Manfield (Wellcome Trust-funded Biomolecular Interactions Facility, Astbury Centre; 062164/Z/00/Z), and Helena Brown (Sorby Environmental Fluid Dynamics Laboratory). For the purpose of Open Access, the authors have applied a CC BY public copyright licence to any Author Accepted Manuscript version arising from this submission.

## Notes

### Competing Interest Statement

The authors have declared no competing interest.

