## Supplementary figures and tables for "Towards Modular Engineering of Cell Signalling: Topographically-Textured Microparticles Induce Osteogenesis via Activation of Canonical Hedgehog Signalling"

***Table S1****: Thermal conditions used in T100 Thermal Cycler (Bio-Rad) for cDNA synthesis.*

| Cycling Step | Temperature | Duration |
|---|---|---|
| Priming | 25°C | 5 minutes |
| Reverse transcription | 42°C | 30 minutes |
| Heat-inactivation | 85°C | 5 minutes |
| Hold | 4°C | ∞ |

***Table S2****: Primer sequences used for quantitative real-time PCR analysis of gene expression.*

| Target Gene | GenBank Accession No. | Direction | Primer Sequence 5'-3' | Concentration (μM) | Annealing Temp (°C) | Amplicon Length(bp) |
|---|---|---|---|---|---|---|
| *Gli1* | AB025922 | Forward | GTCTGACTCCACCGGATTGG | 5 | 60 | 132 |
| | | Reverse | TGAGGATAGGGACCCGACTG | | | |
| *Smo* | NM_176996 | Forward | AACTATCGGTACCGTGCTGG | 5 | 60 | 607 |
| | | Reverse | CATCATGGGAGACAGTGTGC | | | |
| *Patch1* | NM_008957 | Forward | TCAGTTGACTAAACAGCGTCTGGTA | 5 | 60 | 82 |
| | | Reverse | GACCCAAGCGGTCAGGTAGAT | | | |
| *Runx2* | NM_001146038 | Forward | CAGTCCATGCAGGAATATTTAAGGC | 5 | 60 | 135 |
| | | Reverse | AGAAGCTTTGCTGACACGGT | | | |
| *Bglap2* | NM_001032298 | Forward | GGTAGTGAACAGACTCCGGC | 5 | 60 | 177 |
| | | Reverse | GGGCAGCACAGGTCCTAAAT | | | |
| *Gapdh* | GU214026 | Forward | CCATCACCATCTTCCAGGAG | 3 | 55 | 322 |
| | | Reverse | GCATGGACTGTGGTCATGAG | | | |

*Abbreviations: Gli1, glioma associated oncogene homolog 1; Ptch1, Patched1; Smo, Smoothened; Runx2, Runt-related transcription factor2; Bglap2, bone gamma-carboxyglutamate protein 2; Gapdh, Glyceraldehyde-3-Phosphate Dehydrogenase.*

***Table S3****: Thermal conditions used for RT-qPCR.*

| Cycling Step | Temperature | Duration | Cycles |
|---|---|---|---|
| Enzyme activation | 95°C | 30 seconds | 1 |
| Denaturation | 95°C | 5 seconds | 35 |
| Annealing/extension | 55-60°C | 5 seconds | |
| Melting curve | 65-95°C (increment 0.5°C) | 5 seconds/step | 1 |

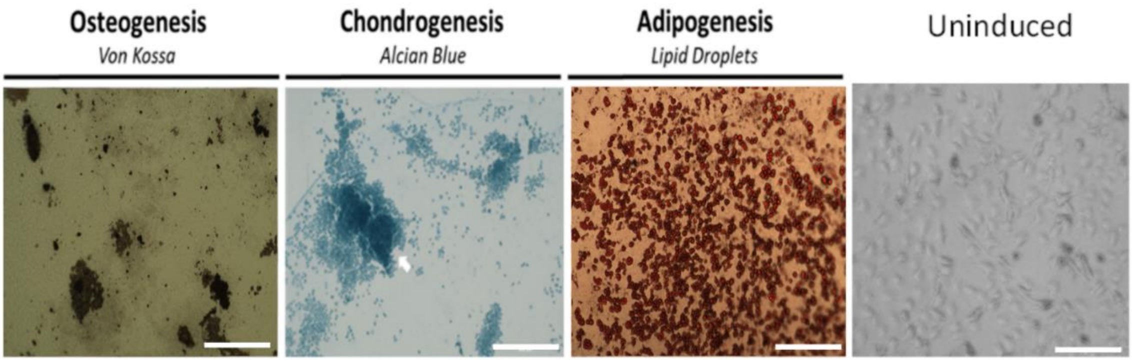


***Figure S1: Tri-lineage differentiation potential of 2D-cultured C3H10T1/2 cells**. To verify the differentiation capacity of C3H10T1/2, biochemically-induced osteogenic (Gibco, UK), chondrogenic (Gibco, UK) and adipogenic (Gibco, UK) differentiation was carried out for 21 days following the manufacturer protocol. Osteogenic differentiation was assessed by Von Kossa staining (Sigma-Aldrich) of the calcified matrix. Cultures showed differentiation down the chondrocyte lineage, as confirmed by Alcian Blue staining pH 1.0 (American Mastertech). Brightfield image showing the formation of lipid droplets and typical adipocyte morphology (124) after staining with Lipid Droplets Assay Oil Red O Solution (Cayman Chemicals). Cells were imaged with EVOS FL Auto 2 microscope (Thermo-Fisher) (Scale bar = 500µm).*

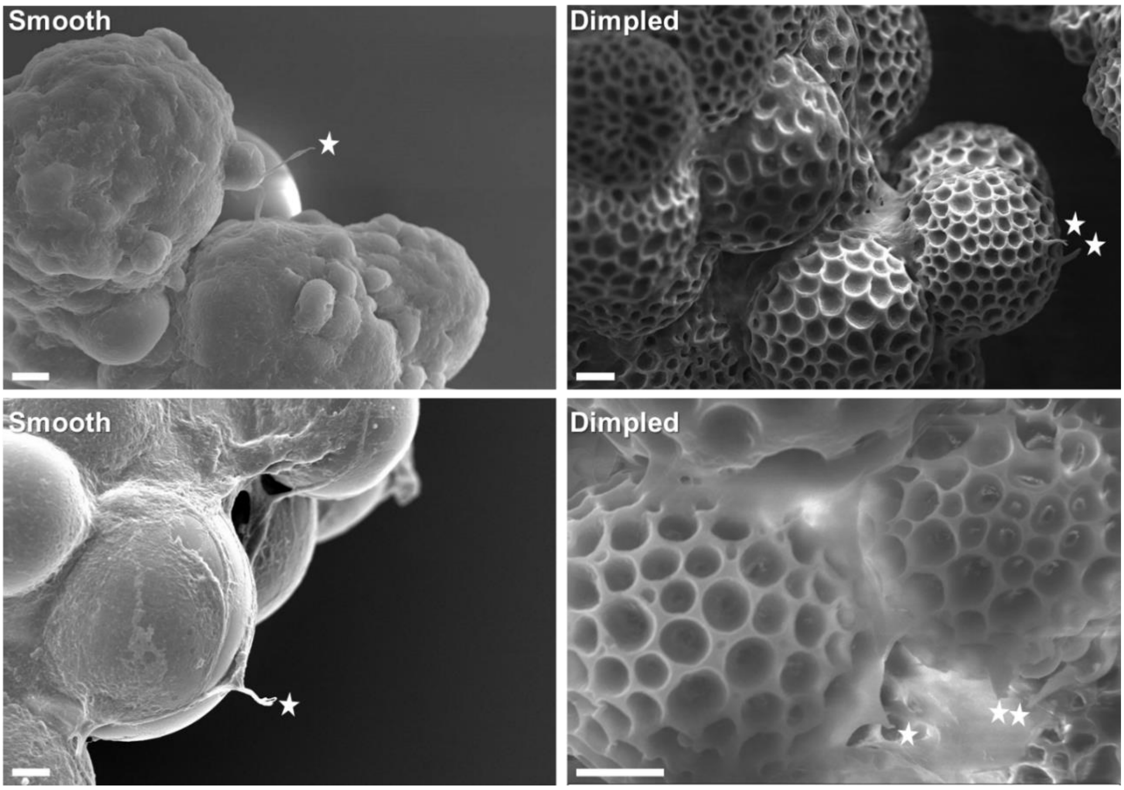


***Figure S2**: Scanning electron microscopy images showing cellular morphologies of C3H10T1/2 cells and the cellular protrusions observed 3 days post-seeding on microparticles. White stars indicate cell protrusions (Scale bar= 10µm).*

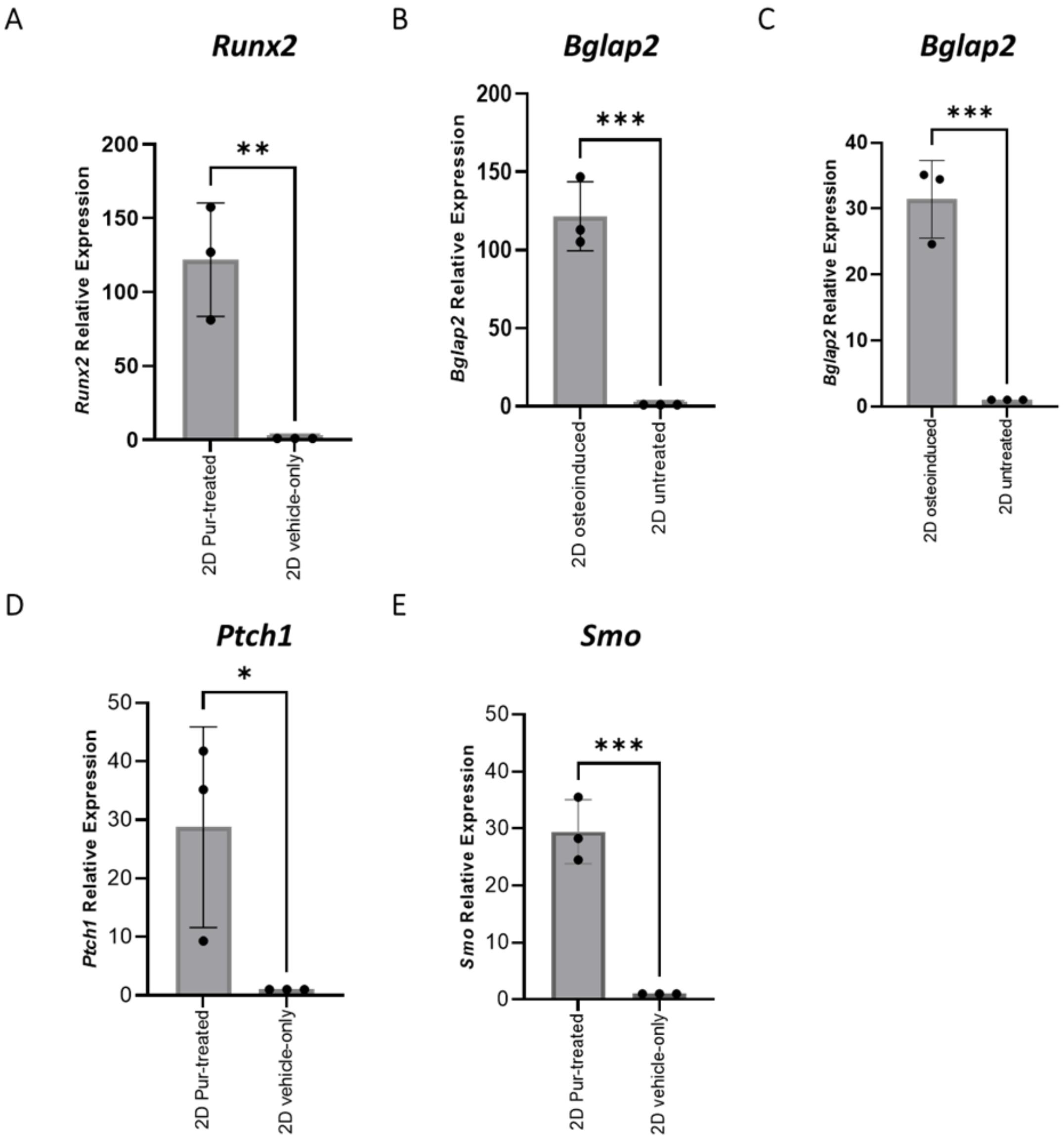


***Figure S3: Expression levels of Hh- and osteogenic-related genes in 2D cultures using biochemical stimulation.*** *Relative qPCR analysis of (A) Runx2, (B) Bglap2 after 3 days and (C) 7 days in culture, (D) Ptch1 and (E) Smo, represented as fold change normalised to GAPDH and relative to either 2D-untreated or 2D vehicle-only controls. C3H10T1/2 cells treated with 2µM purmorphamine were considered as positive controls for the expression of Runx2, Ptch1 and Smo, and expression levels were normalised to 2D vehicle-only controls (containing 0.06% DMSO). C3H10T1/2 cells treated with osteoinductive media were considered as positive (osteo-induced) controls for the expression of Bglap2, and the expression levels were normalised to 2D untreated negative controls. Statistical significance was determined using unpaired Student's t-test. Values are means ± standard deviation for three independent biological repeats. *$p<0.05$, **$p<0.01$, ***$p<0.001$.*

*Abbreviations: Gli1, glioma associated oncogene homolog 1; Ptch1, Patched1; Smo, Smoothened; Runx2, Runt-related transcription factor2; Bglap2, bone gamma-carboxyglutamate protein 2; Gapdh, Glyceraldehyde-3-Phosphate Dehydrogenase; Pur-treated, purmorphamine-treated*

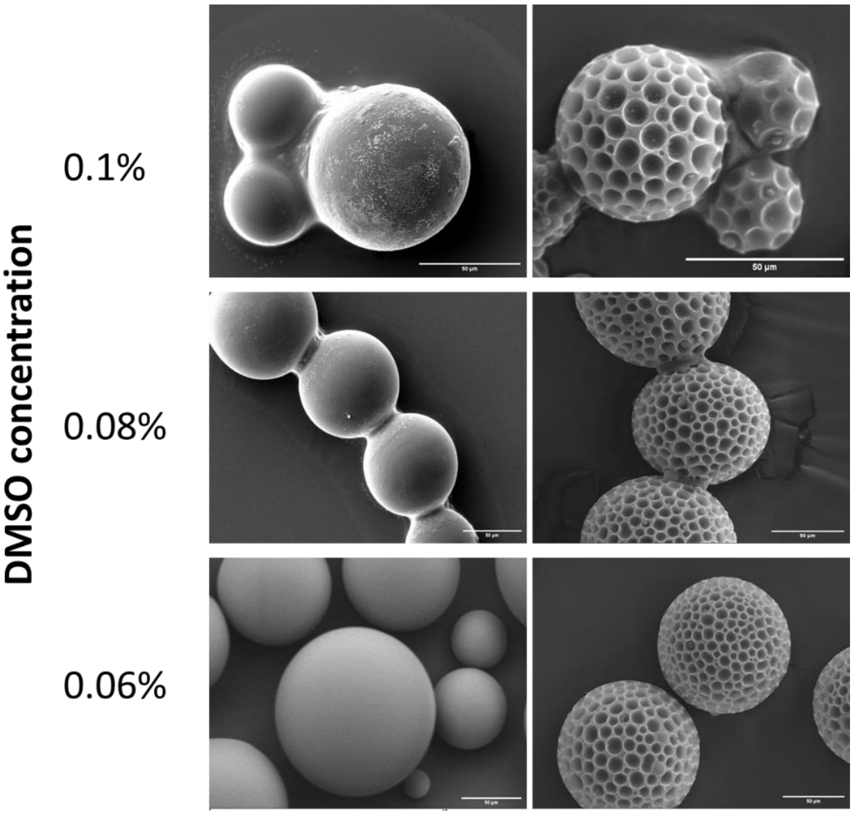


***Figure S4: Optimisation of DMSO concentration to maintain surface topographical features.*** *Representative SEM images showing the effect of three different concentrations of DMSO on the topographical features of the fabricated microparticles. Different DMSO concentrations were prepared by adding 1, 0.8 and 0.6 µL of DMSO in 1mL serum-reduced media to yield 0.1%, 0.8% and 0.06%, respectively. (Scale bar= 50µm).*
